# Distinct phases of immune system programming during ART-suppressed immunodeficiency virus infection

**DOI:** 10.64898/2026.04.29.721702

**Authors:** Maanasa Kaza, Benjamin Varco-Merth, GW McElfresh, Sebastian Benjamin, Gregory J. Boggy, Morgan Chaunzwa, Shana Feltham, Sohita Ojha, Karina Belica, Andrea N. Selseth, Michael Nekorchuk, Kathleen Busman-Sahay, Brandon F. Keele, Dan H. Barouch, Jeffrey D. Lifson, Jacob D. Estes, Scott G. Hansen, Afam Okoye, Louis J. Picker, Benjamin N. Bimber

## Abstract

People living with HIV (PLWH) on suppressive antiretroviral therapy (ART) can face non-AIDS complications, partially driven by chronic immune activation. To define immune perturbations during ART-suppressed viral infection, we performed longitudinal single-cell transcriptomic and plasma proteomic analysis of rhesus macaques infected with SIVmac239M and ART-treated for 70 weeks. We identified broad, bi-phasic immune changes. Acute infection involves an interferon-driven signature, correlated with viral replication, that largely resolves with viral control. Cell-associated virus correlated with interferon-stimulated genes in most tissues; however, this was blunted in gut-associated lymph nodes, a feature that may contribute to reservoir persistence. Separate alterations manifest 54-66 weeks-post-infection, after 40 weeks of viral suppression, including broad TGF-β and NF-kB signaling and discrete bursts of inflammatory monocytes, largely restricted to bone marrow. These data highlight the biphasic remodeling of long-term ART-suppressed HIV, identifying specific tissues and cell populations with dysregulation, with implications for the treatment of PLWH.

## INTRODUCTION

The HIV epidemic remains an important health problem, with more than 40 million PLWH worldwide and an estimated 1.4 million new cases in 2024^1^. ART can suppress viral replication to near-undetectable levels, dramatically improving the prognosis of PLWH and reducing transmission. However, current ART drugs and regimens do not impact already infected cells and do not eradicate HIV^2^. Many lines of evidence highlight the negative impacts and health consequences from persistent HIV infection, even in the setting of effective viral suppression by ART. PLWH face increased risk of many chronic conditions, including diabetes, neurocognitive disorders, cardiovascular disease and certain malignancies^3^. Many non-AIDS comorbidities have been linked to chronic inflammation and immune activation, which have also been suggested to cause premature aging (often termed ‘inflammaging’)^4–8^. Further, many proposed HIV curative therapies rely at least in part on the immune system for clearance of the virus^9^. Therefore, there is a clear need to understand the impact of ART-suppressed HIV infection on the immune system and the overall health of PLWH.

HIV infection has severe and well-documented consequences for the immune system. Uncontrolled infection results in depletion of CD4^+^ T cells, expansion of inflammatory monocytes, increased gut microbial translocation and associated inflammation, and lymphoid tissue fibrosis^6,10–13^. ART-mediated suppression of HIV mitigates many of these effects, resulting in at least partial immune reconstitution^10,14–16^. However, since ART does not completely eradicate HIV-infected cells, immune activation and inflammation persist, which can lead to immune dysregulation^15^. These chronic effects occur even when ART is initiated very early after the onset of infection^17^.

Improved understanding of the immune consequences of ART-suppressed HIV could improve the care of PLWH, including the treatment of comorbidities and the development of proposed HIV curative therapies that rely on the immune system. Controlled non-human primate studies provide a critical complement to human studies and allow the collection of data that is virtually impossible to generate from human cohorts^18^. NHP studies allow for control over the strain, route, and dose of viral infection, as well as the timing of ART initiation and its precise duration at the time of sampling. This enables a more precise evaluation of immune perturbations over time. Extensive tissue sampling, which is only practical in an animal model, is essential because immune measurements from blood do not always correspond to the immune environment of tissues^19^, and lymphoid tissue, especially the gut, is a key reservoir site^20–22^. Because pre-infection samples are readily available, NHPs also allow for precise contrasts between immune cells during stable viral suppression and pre-infection to identify chronic immune modifications that would not be apparent without pre-infection data.

## RESULTS

### RM Cohorts

To investigate the transcriptional and immune dynamics of long-term ART-suppressed SIV infection, we analyzed longitudinal samples from 24 male rhesus macaques (4-6 years old at study initiation) intravenously inoculated with 5000 infectious units of barcoded SIVmac239M, with ART (TDF/FTC/DTG)^23^ starting 9 days post infection (DPI) and maintained for up to 70 weeks post infection (WPI) (**Fig. 1A**). Samples were collected from PBMC and lymphoid tissues, including mesenteric lymph nodes (MesLN), peripheral lymph nodes (PeriLN), bone marrow, and spleen, during acute infection (12 DPI, n=98 samples), early ART suppression (21 WPI, n=98 samples), and late ART suppression (54 and 66 WPI, n=298 samples). Additional PBMCs were collected at 4 WPI (n=23 samples). A total of 626,391 single-cell transcriptomes were analyzed. This RM cohort was previously used to determine the origin sites of post-ART rebound by comparative analysis of barcode-specific viral RNA expression in tissues on-ART and then early off-ART^22^. Our study uses independent transcriptomic and proteomic data and is focused on immune changes elicited by ART-suppressed SIV. To isolate ART-specific effects in the absence of SIV infection, PBMCs were collected from an independent cohort of six male uninfected macaques (5-7 years old) administered the same ART regimen for six weeks (n=6 samples per timepoint, n=8,567 cells) (**Fig. 1B**).

**Figure 1.**
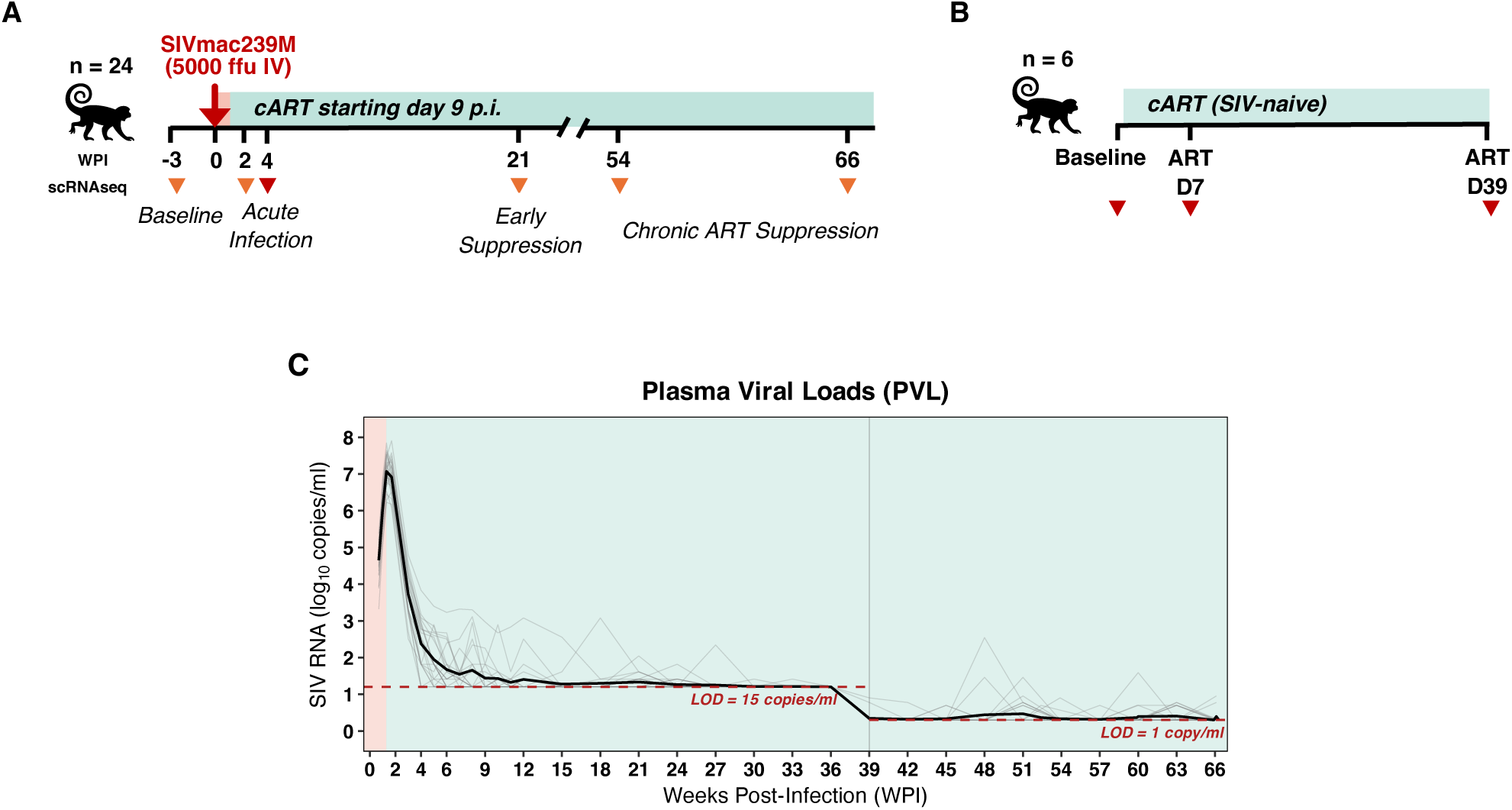
Overview of study design and viral kinetics. **A)** Schematic of the SIV/ART primary cohort (n = 24) showing intravenous SIVmac239M infection (5,000 FFU) followed by ART initiation at 9 days post-infection (DPI) and continued through 66 weeks post-infection (WPI). Samples were collected at predefined timepoints from PBMC, mesenteric lymph nodes (MesLN), peripheral lymph nodes (PeriLN), bone marrow, and spleen; orange triangles); PBMC only were collected at 4 WPI (red triangle). **B)** SIV-naïve comparison cohort (n = 6) treated with ART for 42 days, with PBMC sampled at baseline, ART Day 7 (D7), and ART Day 39 (D39) (red triangles). **C)** Plasma viral load (PVL) trajectories across the primary cohort; individual animals are shown in grey and the cohort median in black. Red dashed line indicates the assay limit of detection (LOD; 15 copies/ml through 36 WPI; 1 copy/ml from 39 WPI onward); blue shading denotes periods on ART.

Longitudinal plasma viral loads (PVL) and tissue cell-associated viral loads (CA-VL) including SIV RNA (CA-vRNA) and SIV DNA (CA-vDNA) from these 24 RM have been previously published^22^, but are reproduced here to enable the integration of transcriptional profiles with systemic and tissue-level viral burden (**Fig. S1**). PVLs peaked at 9-12 DPI and then declined rapidly after ART initiation to less than 15 copies/mL by 15 weeks in almost all subjects. Viral suppression was maintained less than 1 copy/mL in virtually all samples from 39 WPI, with limited low-level viremic blips detected in some samples (**Fig. 1C**). CA-VLs followed a similar pattern, with high CA-vRNA and CA-vDNA at 12 DPI with subsequent decline on ART (**Fig. S1**); CA-vRNA decreased more rapidly than CA-vDNA, as evidenced by the steep decrease in CA-vRNA to CA-vDNA ratio, consistent with prior observations of SIV dynamics under early initiation of suppressive ART^24^.

### Overview of single-cell RNA-seq (scRNA-seq) analysis methodology

The primary data type used in this study is single-cell RNA-seq (scRNA-seq) data, which we collected across the timepoints and tissues indicated above. While this provides single-cell resolution, most analyses were performed using pseudo-bulking, whereby individual cells are grouped by major cell type (T and NK cells, B cells, and myeloid cells), gene expression is aggregated within sample for each cell type, and these pseudo-bulk profiles are used for differential gene expression analyses that are similar to traditional bulk RNA-seq methods^25^. We selected this level of granularity because these coarse cell subsets represent distinct immune lineages and have clearly separable transcriptional profiles when measured by scRNA-seq^26^. The pseudo-bulking approach offers several advantages relative to true single-cell or bulk RNA-seq. Relative to bulk RNA-seq, pseudo-bulking by cell type allows more precise measurement of changes within each cell type and more clearly differentiates changes in gene expression from changes in cell composition. Aggregating single-cell expression profiles into a per-sample profile avoids gene dropout issues present when working with individual cells, and summing expression to the sample level avoids cellular independence assumptions, with significantly improved type-I error rates in differential gene expression analyses compared to typical single-cell methodologies^27–29^. In many cases we employ pseudo-bulking in tandem with single-cell resolution analyses to more precisely map the cellular origin of transcriptional changes identified in the pseudo-bulk analyses.

### Longitudinal Transcriptional Changes in Acute and ART-suppressed SIV Infection

To perform an unbiased evaluation of the major sources of transcriptional variation in our dataset, we first applied dimensionality reduction (PCA/UMAP) and unsupervised clustering to cell type-level pseudo-bulk profiles (**Fig. 2A**). This analysis of high-dimensional transcriptional similarity across samples revealed an unexpected pattern. Acute HIV/SIV infection is characterized by systemic viral replication and drives broad changes in the immune system^30–35^. ART-mediated suppression of viremia reduces this overt immune activation and, in many ways, is generally expected to return cells to a more baseline-like state^36^. However, all pseudo-bulk samples from the late ART-suppressed phase (54-66 WPI), during which viremia has been ART-suppressed to near-undetectable levels for more than 40 weeks, clustered separately from earlier timepoints across all immune cell types (**Fig 2A**). The remaining timepoints, baseline, acute infection (12 DPI), and early ART (21 WPI), clustered more closely together. This does not indicate a lack of changes during acute infection but rather indicates the late ART phase is associated with strong, distinct transcriptional state. The separation between early and late ART samples is also notable given the near-complete viral suppression at both timepoints (**Fig. 1C**), and suggests that early ART samples are transcriptionally closer to the pre-challenge state than late ART samples (**Fig. 2A**). To further illustrate this, we calculated the PCA distance between the pre-infection samples and either acute (12 DPI) or late ART samples (**Fig. 2B**). Across all tissues and cell types, late ART samples were farther from baseline than acute infection samples.

**Figure 2.**
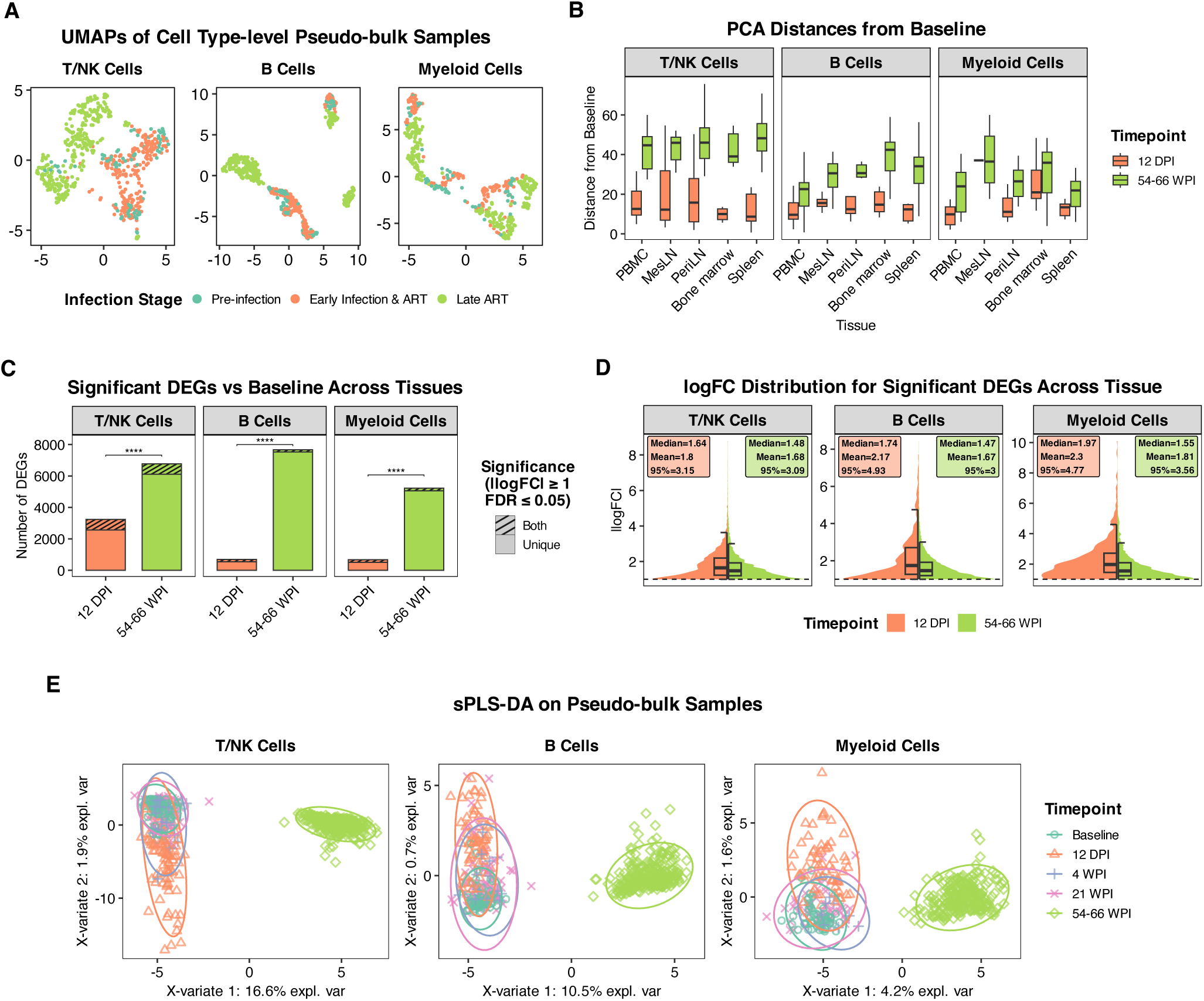
Acute infection and late ART drive distinct, tissue-wide transcriptional shifts across immune compartments. **A)** UMAP embeddings of pseudo-bulk profiles aggregated by sample and immune cell type, colored by infection stage. Late ART (54-66 WPI) samples cluster separately from pre-infection and early infection (12 DPI, 4 WPI, 21 WPI) samples. **B)** Box plots showing the PCA-based distances from each tissue’s baseline profile, comparing acute infection (12 DPI) and late ART (54-66 WPI) across tissues and cell types. **C)** Number of significant differentially expressed genes (DEGs) versus baseline pooled across tissues at 12 DPI and 54-66 WPI (|logFC| ≥ 1, FDR ≤ 0.05), with overlap and timepoint-unique DEGs indicated. Significance tested using a quasi-Poisson GLM of DEG counts adjusted for tissue and cell type, testing 12 DPI vs 54-66 WPI contrasts with FDR correction. **D)** Distributions of absolute logFC for significant DEGs across tissues at 12 DPI and 54-66 WPI, showing higher logFC magnitudes at 12 DPI. **E)** Sparse partial least squares-discriminant analysis (sPLS-DA) of pseudo-bulk samples across longitudinal timepoints within each cell type, highlighting clear multivariate separation of the late ART timepoint (54-66 WPI) along the first two components (variance explained shown on axes).

To validate this finding with complementary computational methods, we performed differential gene expression analysis comparing baseline with either acute infection or late ART and then compared the resulting sets of differentially expressed genes (DEGs). Supporting the PCA/UMAP analysis, significantly more DEGs were identified in the late ART versus baseline comparison than in the acute infection versus baseline comparison (**Fig. 2C, Table S2**). While the number of DEGs was higher at late ART, the absolute log-fold-changes (logFC) among these DEGs tended to be larger during acute phase, suggesting that acute viremia elicits stronger transcriptional perturbations, in a smaller set of genes, that substantially resolve with ART-mediated viral suppression, followed by broader transcriptional remodeling during long-term ART-suppressed SIV that emerges over time (**Fig. 2D, Table S2**). Because statistical significance and false discovery rate estimates can be influenced by sample size, we repeated these analyses using a down-sampled dataset, normalizing the number of samples in each group and obtained comparable results.

As a third approach, we performed sparse partial least squares discriminant analysis (sPLS-DA), a semi-supervised dimensionality reduction method^37^. For each cell type, this analysis identified two components, with one separating the late ART samples from earlier timepoints (**Fig. 2E**, **x-axes)** and another separating the acute phase samples from the remaining timepoints (**Fig. 2E**, **y-axes**). In all cell types, a higher proportion of transcriptional variance was explained by the late ART component (4.2-16.6% of total variance) than the acute phase component (range 0.7-1.9%), further supporting the observations above. When we inspected the gene loadings for these components, we found that the acute phase components for all cell types heavily weighted interferon-stimulated genes (*IFI27*, *OAS1*, and *IFI44*), while the components associated with late ART were more complex, including genes associated with stress response (*TMEM259*) and cell survival (*UBALD2*) (**Fig. S2**). Collectively, these data demonstrate that immune cells during ART-suppressed SIV accumulate changes that are orthogonal to the transcriptional programs elicited during acute viremia, as opposed to the continuation of the same programs at lower magnitude. All major immune cell types accumulate broad transcriptional changes that diverge from baseline states, a pattern that was only apparent because our study design included pre-infection samples.

### ART treatment elicits modest transcriptional changes across immune lineages in the absence of SIV infection

The transcriptional reprogramming identified above could stem from a combination of residual SIV-induced inflammation, effects of ART, or the combination of both. It was therefore essential to separate the effects of ART administration from SIV. We administered the same ART regimen used for the treatment of infected RM to an independent cohort of uninfected RM, collected longitudinal PBMC samples, and performed scRNA-seq. We examined differentially expressed genes at post-ART timepoints (ART D7 and ART D39) relative to baseline (**Fig. S3, Table S3**). While the number of DEGs and magnitude of changes was low, we identified differentially expressed genes in all immune subsets. These changes can be grouped into two categories: transient (increased relative to baseline at ART D7 but resolved by D39) and sustained (increased at D7 and D39). In T/NK cells, we observed a transient induction of lymphocyte survival and homeostasis genes such as *GIMAP1/4/5/8*^38^ and a mild interferon-stimulated gene (ISG) program (*MX1/2*, *OAS2*, *APOL2*) (**Fig. S3A**). NK cells showed an elevation in *CX3CR1* expression at ART D7, which is linked to NK cell maturation and migration^39^. In B cells, the transcriptional profile was consistent with a coordinated endoplasmic reticulum (ER) stress and secretory response with transient increases in *PPIB*, *SEC61B*, *VCP*, and *DERL3* together with the plasma cell linked immunoglobulin assembly factors *MZB1* and *JCHAIN* (**Fig. S3B**). In myeloid cells, we also observed a low-level increase in ISGs (*IFI6*, *MX1*, *EPSTI1*), along with increased immune-regulatory receptors (*CD163*, *CLEC4A*) (**Fig. S3C**), particularly in monocytes and DCs. Collectively, these data demonstrate that ART drugs induce modest transcriptional changes in immune cells, including transient increase in interferon-related genes. These results provide interpretive context for results obtained in ART suppressed SIV infection. While the overall magnitude of these changes is small and the majority resolve within weeks, these data may be relevant to virologic biomarker development.

### Acute SIV infection induces strong, sustained activation across cell types, distinct from ART effects

Next, we examined transcriptional changes elicited during acute SIV infection. We conducted differential gene expression analysis on PBMC samples pseudo-bulked by cell type, contrasting acute phase (12 DPI) with baseline (**Fig. 3 and Table S4**). Genes upregulated during acute phase were clustered according to their longitudinal expression patterns, revealing discrete gene modules with distinct temporal patterns. There was minimal overlap between genes upregulated during acute infection and those induced in the ART-only cohort, indicating that these transcriptional changes are largely in response to SIV infection (**Fig. 3A-B**).

**Figure 3.**
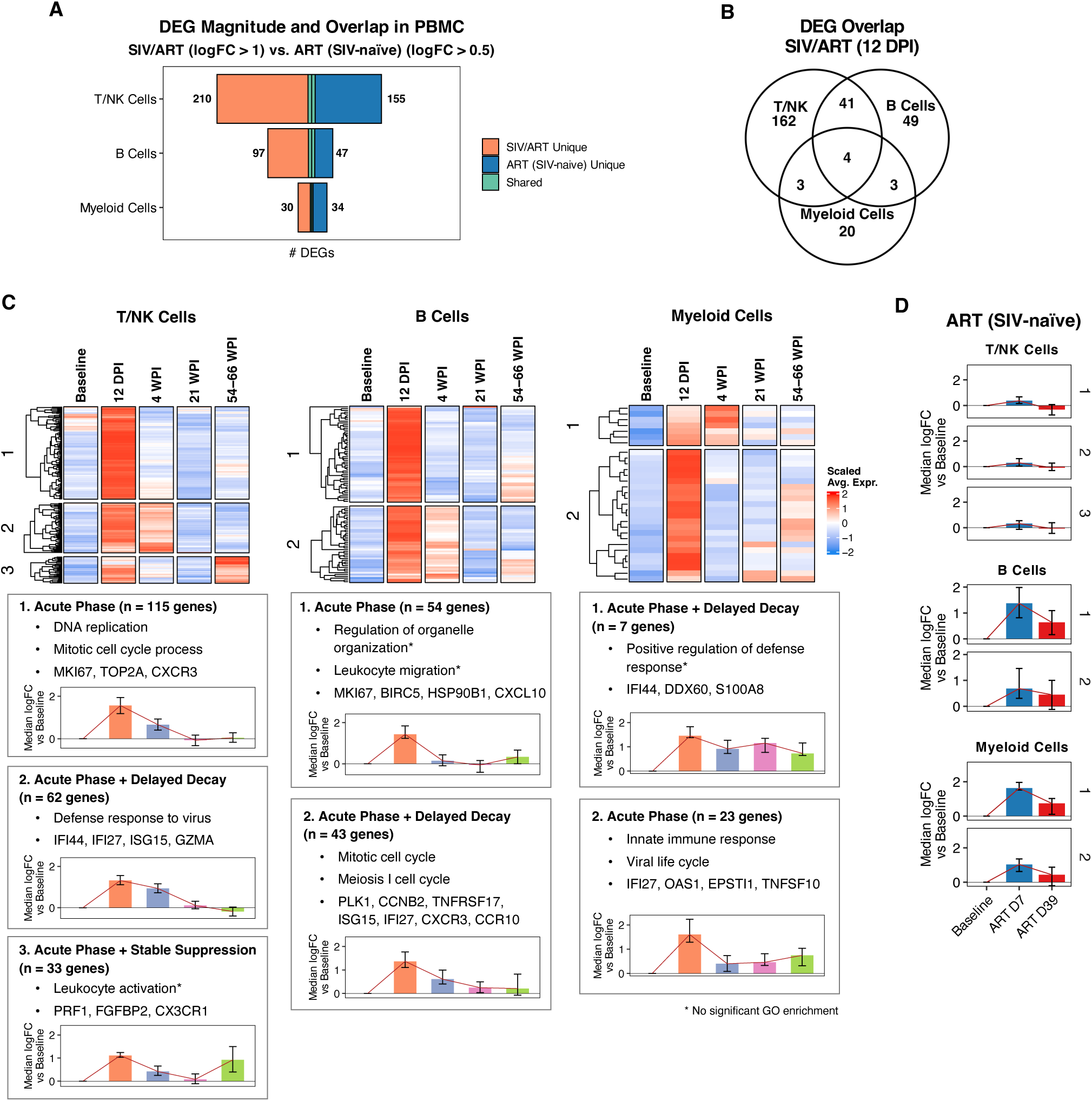
Acute SIV infection elicits strong transcriptional changes across immune subsets. All panels display pseudo-bulked scRNA-seq data from PBMC in the primary SIV/ART cohort, focusing on acute infection (12 DPI). **A)** Bar plots highlight limited overlap of significant DEGs between SIV/ART primary cohort (12 DPI vs baseline; log_2_FC > 1, FDR < 0.05) and ART-treated, SIV-naïve cohort (ART D7 and ART D39 vs baseline; log_2_FC > 0.5, FDR < 0.05) across cell types. **B)** Venn diagram showing overlap of significant DEGs (log_2_FC > 1, FDR < 0.05) at 12 DPI versus baseline across major immune cell types. **C)** Heatmaps of scaled mean expression for significant DEGs at 12 DPI (vs baseline) across longitudinal timepoints in T/NK, B, and myeloid cells. Genes are clustered by expression patterns; representative genes and enriched GO terms are shown for each cluster. Inset bar plots summarize each cluster’s trajectory as the median logFC (vs baseline) over time with IQR. **D)** Bar plots show median logFC vs baseline for the same gene clusters as **(C)** across timepoints the ART-treated cohort.

All three immune subsets (T and NK cells, B cells, and myeloid cells) exhibited gene modules with elevated expression during acute infection (12 DPI), followed by rapid resolution to baseline levels (**Fig. 3C**). While upregulated genes varied between cell types, many shared biological processes were identified. These included cell division and proliferation associated genes (**Fig. 3C**; T/NK module 1 and B cell module 1), consistent with a proliferative burst resulting from generalized immune activation (verified by single-cell resolution analyses in **Fig. S4**). All three cell types also contained gene modules comprised largely of ISGs (e.g., *IFI27*, *IFI6*, *IFI44*, *ISG15*, *MX1*, and *MX2*), which peaked at 12 DPI, decayed with slower kinetics than the proliferation signatures, but returned to near-baseline levels by 21 WPI (**Fig. 3C**; T/NK module 2, B cell module 2, and myeloid module 1). The T/NK module with delayed decay also included cytotoxicity-associated genes (*GNLY*, *GZMA*, *GZMB*, *GZMK*). Single-cell analyses determined that this was explained by an increase in the proportion of cytotoxic T and NK cells at these timepoints (**Fig. S5**). Together, these analyses demonstrate that SIV replication elicits broad immune activation; however, the most overt transcriptional effects are resolved with control of viral replication. Of interest, a subset of these genes was elevated during acute infection, returned to baseline levels, remained low for weeks, and then increased late in the late ART phase (**Fig. 3**, T/NK module 3 and myeloid module 2). This includes an increase in NK cell-associated genes (*PRF1*, *CX3CR1*, *FGFBP2*), which we identified as being caused by an increase in NK cell proportions at both 12 DPI and 66 WPI relative to baseline levels (**Fig. S5A**). In myeloid cells, module 2 was enriched for interferon-stimulated and innate response genes (e.g., *IFI27*, *OAS1*, *EPSTI1*, *SLFN5*, *TNFSF10*), and during late ART we observed re-induction of a subset of this program, including IFN and pattern recognition receptor (PRR)-linked genes (*NLRC5*, *ZBP1*, *TRIM14*, *EPSTI1*) together with activation/remodeling genes such as *SIGLEC1* (CD169), *LGALS8*, and *F13A1*. Similar kinetics for these gene sets were present in all tissues (**Fig. S6**).

### An interferon-stimulated gene signature is correlated with viral burden across tissues

One of the most significant features of acute SIV infection is a broad increase in interferon-stimulated genes. To formalize this signature, we separately calculated the correlation between average expression of each gene differentially expressed, compared to baseline, at 12 DPI in PBMC and both plasma vRNA and cell-associated vRNA (CA-vRNA) at 12 DPI across tissues. The genes with the strongest correlation to viral burden included a conserved set of interferon-stimulated genes (ISGs), containing *ISG15*, *IFI27*, *IFI6*, *OAS2*, *MX1*, *DDX60*, and *USP18* (**Fig. 4A**). To evaluate this signature by sample, we scored cells for enrichment of this ISG module and aggregated enrichment by sample^40^. In PBMC, the kinetics of this score closely mirror PVL, as designed (**Fig. 4B**). When the ART-only cohort was scored using this module, there was a modest and transient increase, which is consistent with our prior findings that ART elicits moderate interferon signaling (**Fig. 4B, right**); however, the magnitude of this was much lower than primary SIV infection (**Fig. 4B, left**). In the SIV-infected cohort, while ISGs decrease significantly as viral replication is controlled by ART, the ISG module remains elevated at 54-66 WPI relative to pre-challenge levels, with statistically significant increases in PBMCs and lymph nodes (**Fig. 4C**). This is consistent with low-level sustained immune activation, despite ART suppression. These data validate the ISG module as a proxy for tissue-specific responses and demonstrate that interferon stimulation of immune cells remains elevated above baseline levels during late ART, despite ART-suppression of viral replication.

**Figure 4.**
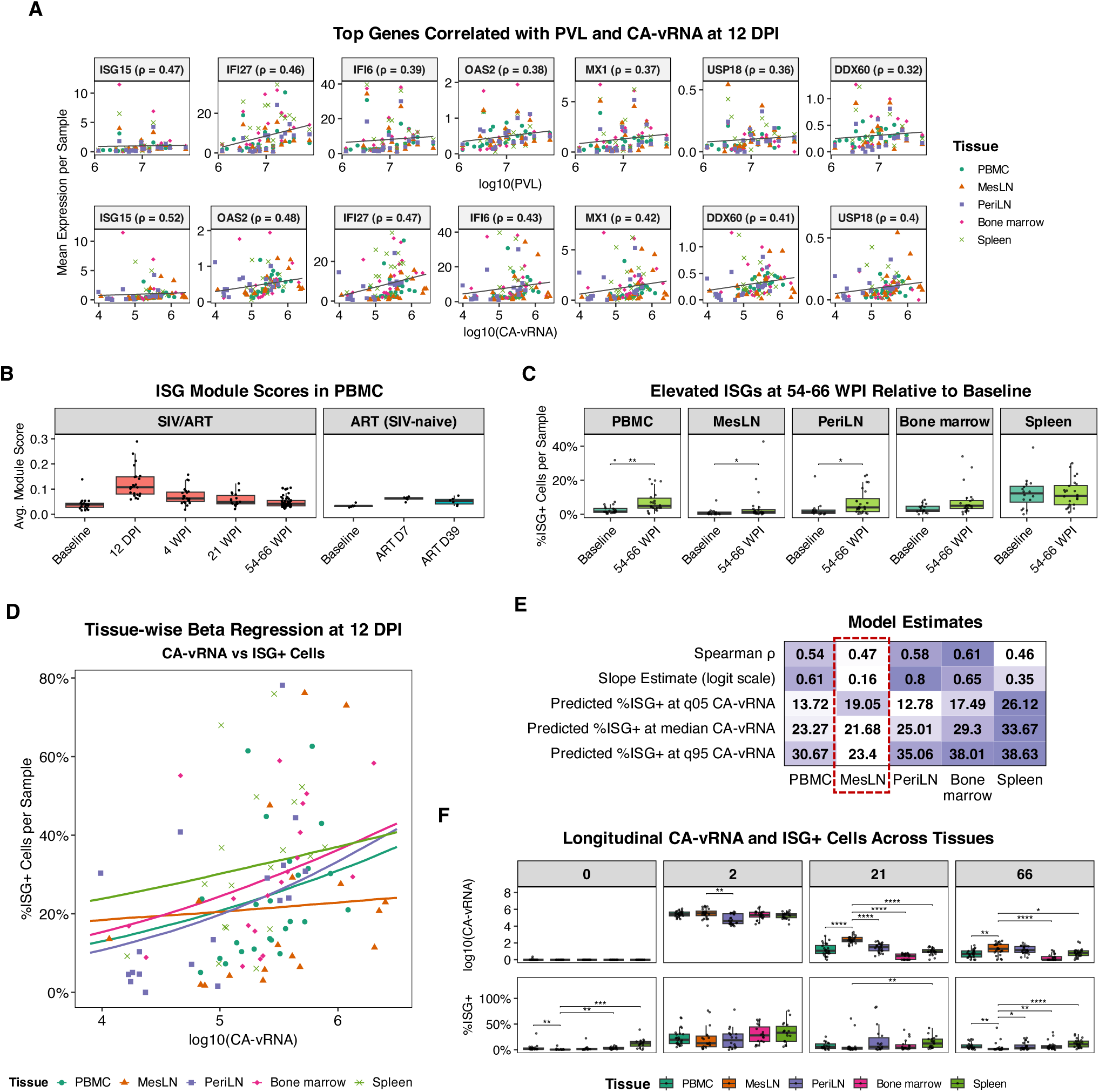
IFN-stimulated gene program tracks viral burden during acute infection, persists through late ART, and reveals tissue-specific differences. **A)** Genes with the strongest associations with viral burden at 12 DPI. Average gene expression per sample is plotted against log_10_(PVL) (top) and log_10_(CA-vRNA) (bottom) across tissues; Spearman ρ is shown for each gene and trend lines indicate linear fits. **B)** Average ISG module scores per sample in PBMC across timepoints for the primary SIV/ART cohort and the ART-treated, SIV-naïve cohort. **C)** Percentage of ISG+ cells per sample at baseline versus late ART (54-66 WPI) across PBMC and tissues, highlighting incomplete return to baseline levels. **D)** Relationship between log_10_(CA-vRNA) and ISG+ cells per sample across tissues, where curves represent tissue-wise beta regression fits. **E)** Summary heatmap reporting Spearman ρ, beta-regression slopes, and predicted ISG+ cells at the 5th, 50th, and 95th percentiles of CA-vRNA for each tissue. **F)** Boxplots of log_10_(CA-vRNA) and ISG+ cells across tissues over time. For both measures, tissues were contrasted with MesLN at each timepoint. Collectively, these data define an ISG-responsive gene signature that tracks viral burden in most tissues and highlight MesLN as a distinct site with higher viral replication without a proportional increase in this pro-inflammatory program.

### Unique immune characteristics of the mesenteric LN

The ISG module provides a tool to explore the correlation between tissue-level cellular responses and local viral burden. We quantified the proportion of ISG+ cells (cells with an ISG module score > 0.2) across tissues and modeled the relationship between ISG+ cells and CA-vRNA at 12 DPI using beta regression. This analysis revealed highly compartmentalized immune dynamics. Bone marrow, peripheral LN, and spleen showed consistent coupling between proportion of ISG+ cells and viral burden, with peripheral LN exhibiting the strongest positive association. In contrast, mesenteric LN displayed a notably flatter slope (**Fig. 4D, orange line**) and consistently lower proportion of model-predicted ISG+ cells at comparable CA-vRNA levels (**Fig. 4E**), indicating that interferon signaling does not scale with viral load in mesenteric LN during acute infection to the same degree as other tissues. This is suggestive of a more immunotolerant environment, which may reflect the need for gut-associated lymphoid tissue to adapt to chronic microbial stimulation. Longitudinal comparisons of CA-vRNA and ISG+ cells across tissues further highlighted unique characteristics of mesenteric LN. ISG+ cells are significantly lower in mesenteric LN relative to PBMCs prior to infection (**Fig. 4F**). While percentage of ISG+ cells increase in all tissues during acute infection, these cells declined sharply in mesenteric LN relative to other tissues by 21 WPI, despite significantly higher CA-vRNA levels. Together, these findings point to a unique immune environment in mesenteric LN that limits interferon responsiveness despite ongoing or residual viral replication, potentially creating a more permissive environment for viral persistence. These characteristics could contribute to both the establishment and maintenance of the viral reservoir in gut-associated lymphoid tissue.

### Longitudinal changes in CD4^+^ and CD8^+^ T cells

Next, we inspected T cells for markers of immune activation. We scored CD4^+^ and CD8^+^ T cells, for gene programs associated with TCR stimulation and cytokine production (**Fig. S7A-B**). While both CD4^+^ and CD8^+^ T cell activation increased during acute infection (12 DPI) in most tissues, this declined by 21 WPI and returned to near-baseline levels in most tissues, with the exception of lymph nodes, which had a small but statistically significant increase in T cell activation at 54-66 WPI relative to pre-infection (**Fig. S7A**). Combined with prior findings, this demonstrates that the mesenteric lymph node has both a blunted interferon response to the virus and an increase in activated CD4^+^ T cell targets.

There were also shifts in key CD4^+^ and CD8^+^ T cell lineage and differentiation markers over the course of infection (**Fig. S7C-E)**. Inhibitory molecules *TIGIT* and *LAG3* were upregulated in CD8^+^ T cells during acute infection but returned to pre-infection levels by 54-66 WPI. A notable longitudinal change was the increased expression of both T-bet (*TBX21*) and *EOMES* in CD8^+^ T cells by 54-66 WPI. Single-cell analyses showed that this was driven by expansion of two distinct populations: EOMES-Hi / T-bet-Lo, and EOMES-Lo / T-bet-Hi, both of which increase over time (**Fig. S7C-D**). These phenotypes have been reported in natural aging and may reflect a similar process induced by SIV infection^41^. Longitudinal changes in CD4^+^ T cells were less pronounced than CD8^+^ T cells and included an acute phase increase of *LAG3*. Expression of T-bet increased during the acute phase, returned to baseline, and then increased again during late ART (54-66 WPI).

### Late-stage monocyte activation and dysregulation in bone marrow

One of the most significant changes during the late ART phase was an increase in myeloid cells expressing pro-inflammatory cytokines (**Fig. 5**). This population was first detected in bone marrow at 54 WPI, although their frequency varied widely between subjects and samples (**Fig. 5A**). These cells are primarily monocytes and express high levels of pro-inflammatory genes, such as *IL1B*, *TNF*, and *CXCL8* (IL-8), and chemokines such as *CCL3* (MIP-1α) and *CCL4* (MIP-1β). This is clearly distinct from the program induced in myeloid cells during acute SIV infection, which is primarily characterized by an increase in interferon-stimulated genes, *CCL5* (RANTES), and the pro-apoptotic cytokine *TNFSF10* (TRAIL) (**Fig. 5B**). In addition to considerable variability between subjects, the frequency of these monocytes also fluctuated within each subject between 54 and 66 WPI, suggesting that in most cases they represent transient blips rather than a stable population (**Fig. 5C**). Note that, while most RM were taken off ART at 70 WPI to investigate viral rebound in our companion study^22^, six RM remained on ART, allowing an additional timepoint to be included in this analysis. Further, the frequency of these cells in the bone marrow was not predictive of their frequency in PBMCs (**Fig. 5D**). Because the 54 and 66 WPI bone marrow biopsies were performed at different anatomical sites, it is also possible that there is compartmentalization of inflammatory monocytes within the bone marrow, rather than fluctuation in their levels over time. Together, these data identify a distinct subset of monocytes that emerges during late ART, is largely restricted to bone marrow, and varies in magnitude between samples. Other studies have identified similar monocyte populations associated with HIV rebound, raising the possibility that these cells might reflect local viral dynamics^42^. However, the frequency of this monocyte population did not correlate with available virologic measurements, including CA-vRNA and CA-vDNA.

**Figure 5.**
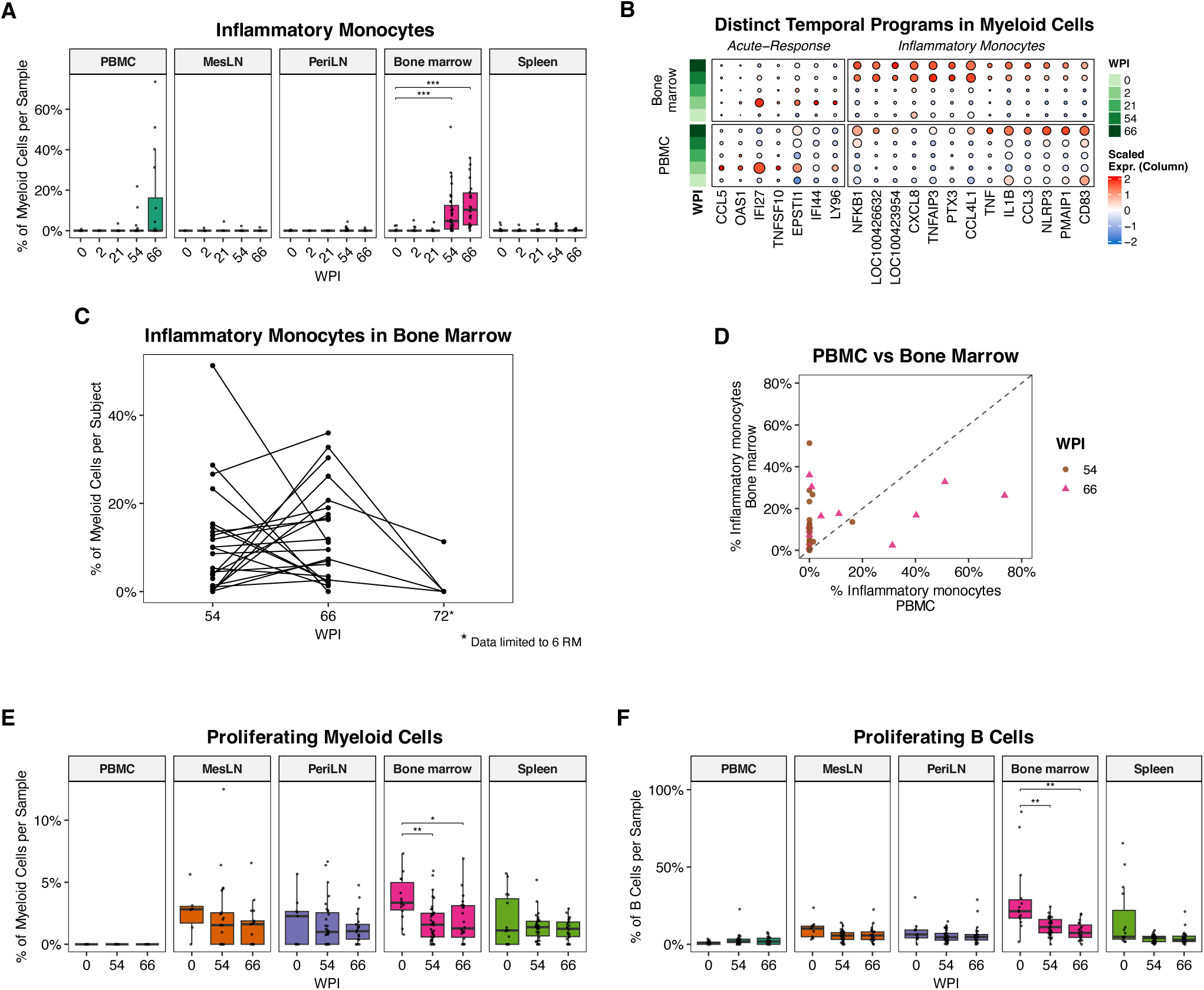
Persistent dysregulation in bone marrow during late ART suppression. **A)** Proportions of inflammatory monocytes across tissues over time, highlighting enrichment during late ART, most prominently in bone marrow and PBMC. **B)** Dot plot of scaled mean expression illustrating that the inflammatory monocyte signature is distinct from the acute infection response in myeloid cells, displayed separately for bone marrow and PBMC across timepoints. **C)** Subject-level trajectories of inflammatory monocyte frequencies in bone marrow across timepoints. The 72 WPI timepoint is only shown for 6 RM, since the other RM were taken off ART at this timepoint. **D)** Scatter plot comparing inflammatory monocyte frequencies between PBMC and bone marrow within subjects at 54 and 66 WPI. **E-F)** Percentages of proliferating myeloid cells **(E)** and B cells **(F)** at the indicated timepoints. For both subsets, there is a significant reduction in proliferation in bone marrow at 54-66 WPI.

Interestingly, we also detected a significant decrease in both myeloid and B cell proliferation in bone marrow at 54-66 WPI relative to pre-infection, potentially reflecting the effects of chronic inflammation (**Fig. 5E-F**). Collectively, these data suggest that the bone marrow is one of the earliest tissues to acquire an overtly dysregulated phenotype and further emphasize the distinct, significant immune alterations that emerge during the late ART phase. Because these inflammatory monocytes can accumulate in bone marrow before they become observable in the blood, they may not always be apparent when sampling blood alone. Even if these cells are primarily restricted to bone marrow, they upregulate several soluble factors that could have systemic effects.

### Altered plasma proteome during ART-suppressed SIV

To complement the transcriptomic analyses, we used the NULISA platform to quantify 119 inflammatory and immune-related plasma proteins at baseline, acute infection, and late ART timepoints (**Table S6**). Analyses were restricted to targets with greater than 50% detectability within the panel (119 of 200; **Table S6B**), as described in the Methods. Principal component analysis separated baseline, acute infection, and late ART samples along the first two principal components (**Fig. 6A**). This indicates that, while acute infection drives the expected strong changes to the plasma proteome, there is a distinct inflammatory plasma protein signature specific to the late ART phase. This pattern is concordant with our transcriptomic data (**Fig. 2**).

**Figure 6.**
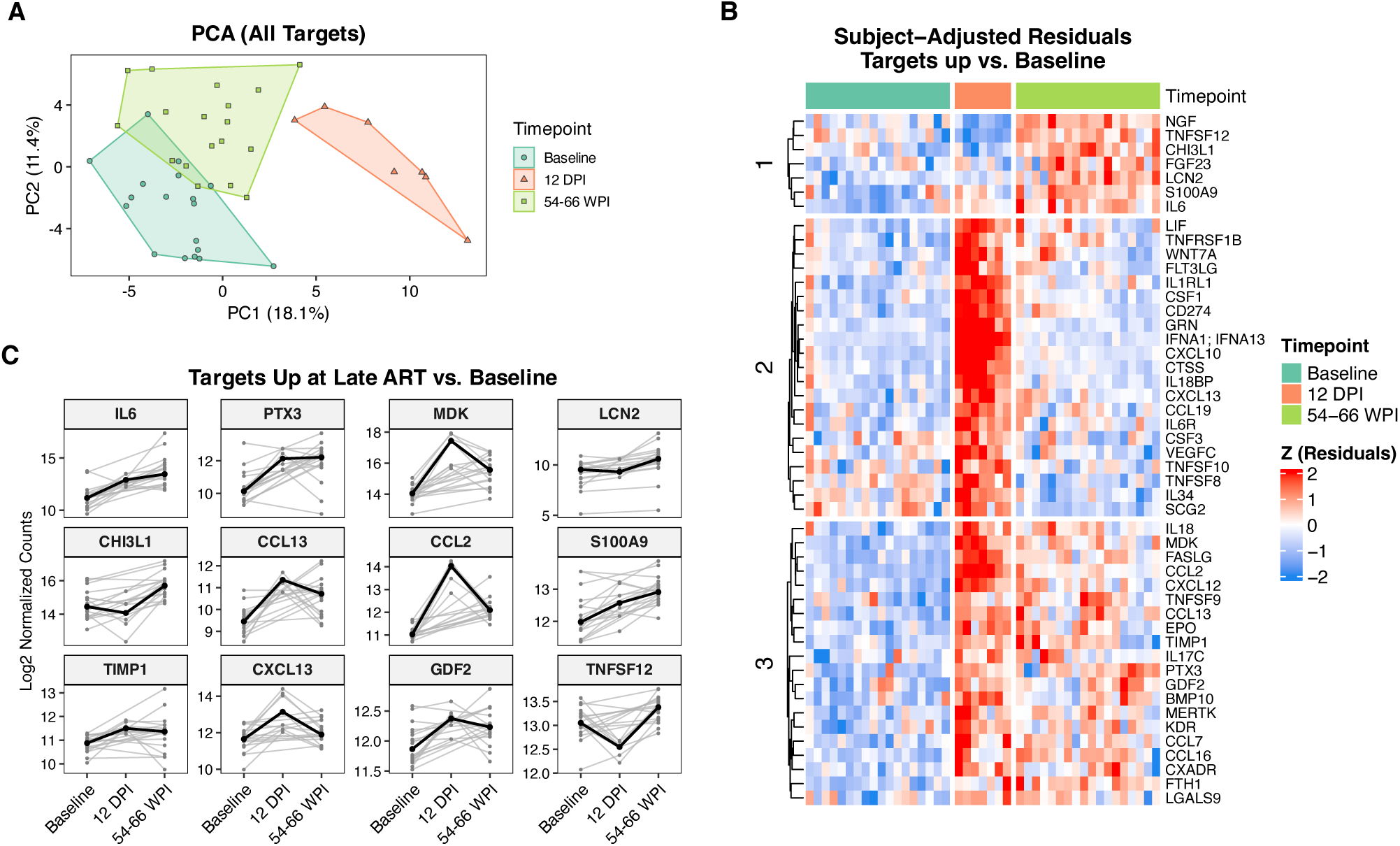
Plasma proteome dynamics across acute SIV infection and long-term ART suppression. **A)** PCA of log_₂_-normalized NULISA counts for 119 inflammatory and immune-related plasma proteins at baseline, acute infection, and late ART. **B)** Heatmap of subject-adjusted residuals (scaled within protein) from mixed-effects models for proteins identified as upregulated relative to baseline at acute infection and/or late ART. Proteins are ordered by hierarchical clustering into three major clusters, highlighting proteins that peak during acute infection versus those that remain elevated or increase further at late ART. **C)** Per-subject trajectories (gray lines) and group medians (black line) for representative proteins significantly upregulated at late ART compared with baseline, shown as log_₂_-normalized counts across baseline, acute infection, and late ART.

To identify plasma proteins that increased from baseline during acute infection and/or late ART we performed mixed-effects modeling and resolved clusters of proteins that either peaked at acute infection and then declined or remained elevated, or increased further, during late ART (**Fig. 6B**). As expected, the proteins that peaked during acute infection included antiviral and cytotoxic mediators such as IFNA1/IFNA13, the interferon inducible chemokine CXCL10, the myeloid growth factors CSF1 and CSF3, and the TNF superfamily ligands TNFSF10 (TRAIL) and TNFSF8, consistent with a transient burst of antiviral and cytotoxic signaling during early infection. These markers returned to baseline levels after ART-mediated viral suppression.

The remaining plasma proteins fell into two categories. The first group was strongly induced during acute infection and remained elevated above pre-infection levels, which included the inflammatory cytokine IL-6, soluble pattern recognition receptor PTX3, chemokines CCL2, CCL13, and CXCL13, and GDF2, MDK, and TIMP1 (remodeling factors). Increased plasma IL-6 and PTX3 are consistent with the scRNA-seq data, which showed elevated expression of these genes in a subset of bone marrow monocytes at 54-66 WPI (**Fig. 5B** and **Table S2**). In contrast, LCN2 (NGAL), CHI3L1, and FGF23, and TNFSF12 (TWEAK) showed relatively modest induction during acute infection but significantly elevated during the late ART phase (**Fig. 6C**). LCN2 can be induced by inflammatory cytokines, and injury of epithelial cells in the intestine, stomach, liver, or lungs during infection^43^. Elevated plasma TNFSF12 is concordant with our scRNA-seq data, which shows elevated expression of *TNFSF12* at 54-66 WPI in NK cells, monocytes, and DCs. TWEAK/Fn14 signaling promotes chronic inflammation and fibrosis and induces pro-inflammatory cytokines, including IL-6 and CCL2^44^. Together, these findings complement our scRNA-seq data and indicate that, in addition to the peak inflammatory response during acute infection, ART-suppressed SIV infection elicits a distinct set of proinflammatory and immunomodulatory changes, including persistent elevation of classical inflammatory mediators and a delayed inflammatory axis associated with neutrophil activation and tissue fibrosis.

### Broad transcriptional dysregulation of immune cells after sustained ART suppressed SIV infection, despite nearly undetectable viremia

Our data demonstrate that the immune cell transcriptomic and plasma proteomic environment after prolonged ART-suppressed SIV infection does not return to a pre-infection state. We identified multiple immune subpopulations with activated/inflammatory states; however, these are discrete populations, often restricted to specific tissues, and represent a small fraction of total immune cells. Our initial PCA/UMAP and differential gene expression analyses identified broad-based changes across all immune cells during the late ART phase, which were absent from earlier timepoints (**Fig. 2A-C**). To define gene programs underlying these broad perturbations, we performed differential gene expression analysis to identify genes upregulated during late ART (54-66 WPI) relative to pre-infection, acute infection (12 DPI), and early ART (21 WPI) in PBMCs (**Table S5**).

These genes were clustered according to their longitudinal expression patterns, revealing cell type-specific programs (**Fig. 7A**; clusters 1-3) and a shared module (**Fig. 7A**; cluster 4). The shared module prominently included *TGFB1* and *SMAD4* and was enriched for the “regulation of TGF-β receptor signaling” pathway. TGF-β1 is a potent immunosuppressive cytokine that restrains T-cell activation and proliferation and has been implicated in promoting HIV/SIV latency and lymphoid tissue fibrosis^13^. We also observed coordinated upregulation of other transcriptional regulators such as *JUND* (AP-1 family transcription factor) and *MAP2K2* (MEK2) (**Fig. 7A**). AP-1 and MAPK/ERK signaling are suggested as drivers of HIV LTR activity, latency establishment, and reactivation^45–47^. Many of the cell type-specific changes are concordant with data from prior figures, including the increase in TBX21 (T-bet) in T and NK cells (**Fig. 7A**, module 1, and **Fig. S7C**) and increases in multiple pro-inflammatory cytokines and chemokines in myeloid cells, including *IL1B*, *TNF*, *CCL3* (MIP-1α), and *CCL4* (MIP-1β) (**Fig. 7A**, module 3, and **Fig. 5**).

**Figure 7.**
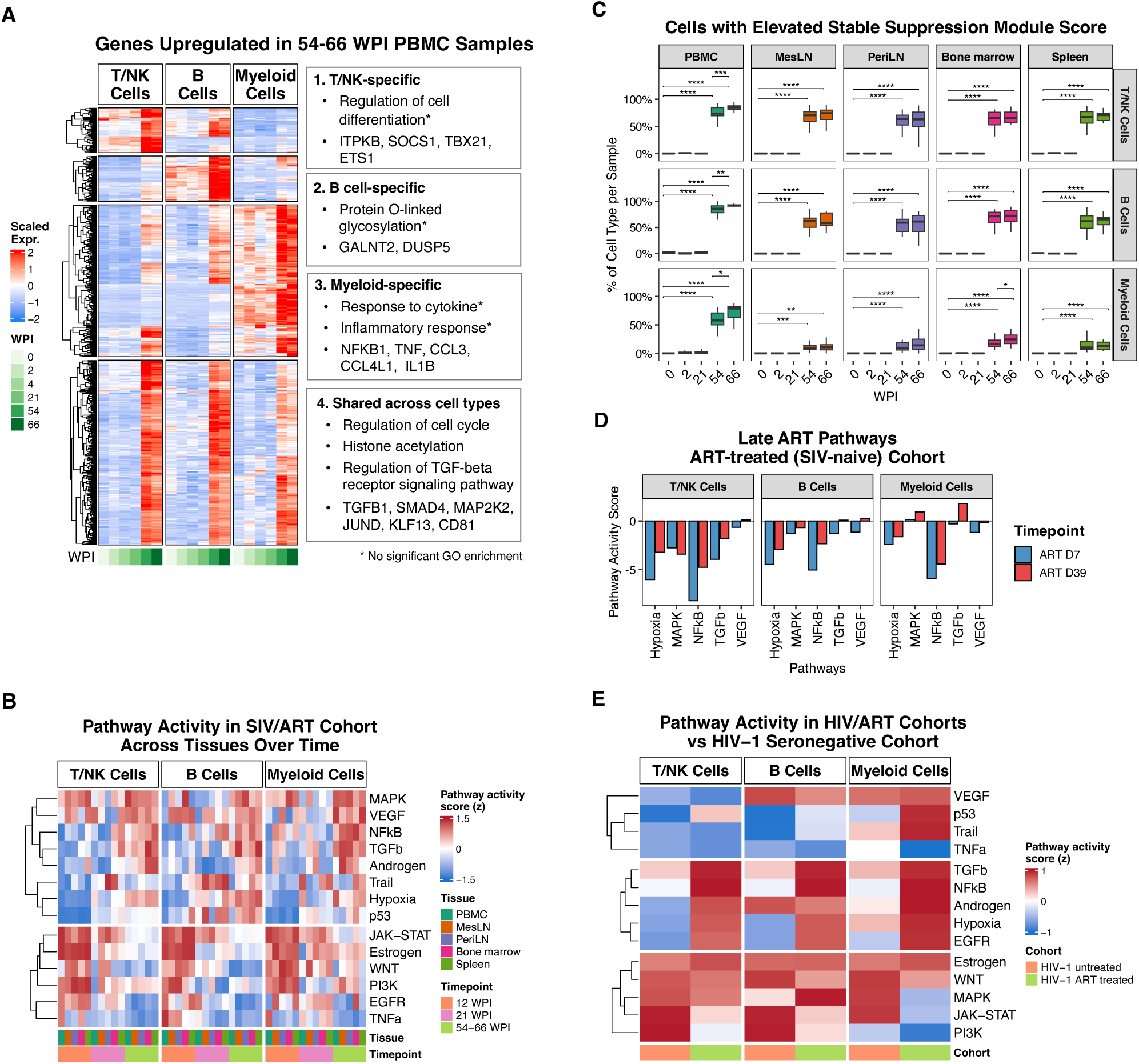
Broad transcriptional remodeling of immune cells after long-term ART-suppressed SIV infection. **A)** Clustered heatmap of genes upregulated during long term ART timepoints relative to pre-infection and early infection/ART, with representative genes and enriched GO terms annotated for each gene cluster. **B)** Pathway activity inference using decoupleR/PROGENy, highlighting distinct programs induced during acute infection and late ART, including increased hypoxia, TGFβ, NF-kB, VEGF, and MAPK signaling at 54-66 WPI. (**C**)Unscaled pathway activity scores for the late ART pathways inferred using decoupleR/PROGENy in the ART-treated, SIV-naïve cohort. Bar plots show inferred activity for hypoxia, MAPK, NF-κB, TGF-β, and VEGF across T/NK cells, B cells, and myeloid cells at ART Day 7 (D7) and ART Day 39 (D39). **D)** Pathway activity inference using decoupleR/PROGENy in HIV-1 untreated and HIV-1 ART-treated cohorts relative to HIV-1 seronegative controls. The untreated HIV-1 cohort showed enrichment of JAK-STAT, estrogen, and PI3K signaling, resembling acute and early infection/ART timepoints in the SIV/ART cohort, whereas the ART-treated HIV-1 cohort aligned with late-ART SIV samples, with increased TGF-β, NF-κB, hypoxia, and androgen signaling.

To complement these analyses, we performed pathway activity inference using the PROGENy database^48^. This analysis further supported the finding that acute SIV infection and sustained ART suppressed SIV infection are characterized by distinct, orthogonal programs. This initial response to SIV infection is consistent with a direct antiviral response, including elevation of the signaling pathways for JAK-STAT (which is stimulated by interferon), WNT, and PI3K (**Fig. 7B**, bottom). These pathways peak during acute infection, returning to near baseline levels by 21 WPI. By 54 WPI, all RM switched to an alternate program that includes elevated TGFβ, MAPK, NF-kB, hypoxia, androgen, and VEGF signaling (**Fig. 7B**, top). The strongest elevation of these pathways occurred in lymphoid tissues, with much more modest changes in PBMCs. To evaluate the breadth of these changes and summarize this effect across tissues, we scored cells for enrichment of a diagnostic late ART gene module based on the genes upregulated across cell types (**Table S7**). In the SIV-infected, ART-treated cohort, the baseline, acute phase, and early ART timepoints across all tissues showed virtually no cells with elevation of this module (**Fig. 7C**). In contrast, at 54-66 WPI, >70% of T/NK cells and >67% of B cells were elevated for this gene program, emphasizing the breadth of its impact (**Fig. 7C**). While enrichment for this module was lower in myeloid cells, this is a technical artifact. Monocytes, which are the primary myeloid cell type in PBMCs, showed comparable levels of enrichment to T and B cells (**Fig. 7C and S9A**). Macrophages, the dominant myeloid cell type in lymphoid tissues, generally had lower module scores. Macrophages also have lower overall RNA expression, which will negatively impact the enrichment score. When we inspected expression of the individual genes in this module, most increased at 54-66 WPI relative to baseline, demonstrating that macrophages upregulate the same program (**Fig. S9B**). The ART-only, SIV-naïve cohort lacked the pathway upregulation observed in late SIV/ART and had minimal expression of the diagnostic module (**Fig. 7D and Fig. S9A**).

To test whether similar immune re-programming is present in PLWH while on ART, we obtained publicly available scRNA-seq data from a human cohort that includes HIV-negative, untreated HIV-positive, and ART-treated HIV-positive subjects^49^. Individuals were mostly male and of African descent. The ART-treated group received variable drug combinations for 27-35 weeks and had PVLs < 400 copies/mL^49^. The untreated HIV-1 cohort was enriched for JAK-STAT, estrogen, and PI3K, resembling the acute phase and early ART SIV samples (**Fig. 7E**). The HIV-1/ART-treated cohort resembled the late ART SIV samples, with increased TGFβ, NFκB, hypoxia, and androgen signaling. Additionally, we identified a monocyte cluster in the human cohort expressing genes similar to the inflammatory monocyte program in the SIV/ART cohort, with shared DEGs showing concordant upregulation across cohorts (**Fig. S9C**). This monocyte population was not evident in the seronegative or HIV-1/untreated groups and was enriched in only 1 of 3 HIV-1/ART-treated subjects, indicating heterogeneity within the human cohort. Together, these data suggest that similar processes occur in ART-suppressed HIV infection and the ART-suppressed SIV cohort.

## DISCUSSION

ART therapy dramatically improves life expectancy and quality of life for people living with HIV. Even if the virus is suppressed to levels undetectable in the plasma, it is not eradicated, and low-level viral activity and immune stimulation persist^2,5,50–52^. Understanding the immune consequences of this is critical for patient care, comorbidities, and proposed HIV cure therapies that depend on the immune system. Because HIV persists primarily in lymphoid tissues, including sites that are difficult to sample in human subjects, controlled non-human primate models of SIV infection can provide critical insight not feasibly gained from human cohorts. Here we present a large and comprehensive dataset to longitudinally characterize immune cell modifications in SIV-infected, ART-suppressed rhesus macaques. One of the primary contributions of this study is breadth. Rather than focus on specific subsets, we employed scRNA-seq to characterize transcriptional and compositional changes to major immune subsets from blood and lymphoid tissues over 72 weeks. We contrasted these cellular measurements against virologic measurements and plasma proteomics to gain insight into the immune response to immunodeficiency virus infection and the alterations to the immune landscape during ART-suppressed infection.

Our data demonstrate that the impact of SIV infection and ART-suppressed SIV can be separated into two phases, characterized by distinct immune cell alterations. Acute SIV infection results in broad immune activation across all tissues, with increased expression of interferon-stimulated genes and elevation of pro-inflammatory cytokines in plasma. This is well documented in both HIV and SIV^4,6^. At the pathway level, we show that acute SIV infection results in transient increases in TNFa, JAK-STAT, WNT, and PI3K signaling. At a cellular level, this corresponds to increased proliferation in T and B cells and increased numbers of cytotoxic T and NK cells. These immune perturbations resolve to near-baseline levels after ART initiation, although the kinetics vary by tissue. However, features like interferon-stimulated gene expression remained slightly elevated relative to pre-infection levels at 66 WPI following more than 30 weeks of complete viral suppression. The sustained, low-level elevation of pro-inflammatory genes that we detect in cellular transcriptomes was also observed in plasma proteomic data. Together, these underscore the fact that even when viremia is controlled to near-undetectable levels, gene and protein programs that are directly connected to the antiviral response remain above baseline.

One of the more surprising findings was the magnitude of the immune changes acquired after sustained viral suppression and the differences between early and late ART phases. Rather than eliciting a gradual return to a pre-infection state, ART suppression resulted in a transient return to baseline, followed by a delayed and broad set of transcriptional and proteomic changes that are distinct from the gene programs correlated directly with viral burden. The late ART programs were dominated by TGF-β signaling, a complex cytokine often associated with immunosuppression, along with increases in NF-κB, VEGF, and androgen signaling. The patterns we observed in RM SIV/ART samples were also identified by reanalysis of a public HIV/ART dataset, suggesting that the RM data have relevance to PLWH^49^. While we did not perform functional assays, these changes are consistent with descriptions of HIV-induced ‘inflammaging’, a state characterized by an imbalance between pro- and anti-inflammatory signals that results in degraded immune function^53^.

Important differences exist between immune cells in blood and local tissue environments. Our findings underscore these differences and identify two new features with potential importance to the care of PLWH. A growing body of literature implicates gut-associated lymphoid tissue as an important site of both viral persistence and early viral reactivation^21,22,54,55^. Indeed, in the same RM used in this study, we documented that gut-associated lymph nodes harbor a larger and more active viral reservoir than peripheral lymph nodes and was a dominant site of initial post-ART rebound. Here, we identified differences in the local antiviral response (interferon-stimulated genes), specific to gut-associated lymphoid tissue. In PBMCs, spleen, bone marrow, and peripheral lymph nodes we show that the level of interferon response scales with increasing CA-vRNA. This relationship is blunted in the gut-associated lymph nodes, indicating that higher viral replication is tolerated without a commensurate increase in interferon signaling and cell-intrinsic immunity. Further, there is a progressive increase in activated CD4^+^ and CD8^+^ T cells in mesenteric lymph nodes throughout the ART-suppressed phase, resulting in increased targets for the virus. These provide a potential mechanism to explain enhanced viral persistence and identify a potential point of therapeutic intervention. Second, we identified cellular dysregulation in the bone marrow that only emerges in the late ART phase. We detected a population of pro-inflammatory monocytes that was sporadically detected in late ART (54-66 WPI) but absent in the early ART phase (21 WPI). Because they are primarily restricted to bone marrow, a tissue less commonly sampled in PLWH, they may be missed in many studies. Even if these cells reflect a secondary consequence of viral-induced inflammation, they express many soluble factors that are increased in plasma, including IL-6, and thus may have an outsized contribution to systemic immune cell reprogramming. While our data did not identify a correlation between local viral levels and these blips in inflammatory monocytes, other studies have directly linked monocyte expansion and inflammatory phenotypes to viral replication, and it is possible there is a less direct connection^17^. Characterizing and reducing the stimuli that drive these cells may reduce the secondary effects associated with ART-suppressed HIV and improve patient care.

There are limitations to this study. All rhesus macaques used in this study were male, and thus we are unable to study sex-specific differences. While our data included a control cohort that received ART alone, this cohort was not monitored for the same duration as the SIV/ART cohort, and thus it is possible that ART drugs contribute to at least some of the immune changes observed during the late ART phase. Additional studies are required to fully separate the impact of ART relative to ART suppressed SIV infection.

This study represents a broad and comprehensive window into the immune cell transcriptional and plasma proteomic changes that occur over the course of SIV infection and ART suppression. While our data broadly align with published features of HIV/ART, this study adds considerable cellular and mechanistic insight, particularly with respect to differences between immunologic measurements of blood versus lymphoid tissues and the kinetics of immune modifications while on ART. These data can inform the development of HIV cures and the treatment of PLWH.

## Resource Availability

### Lead contact

Requests for further information and resources should be directed to and will be fulfilled by the lead contact, Dr. Benjamin Bimber.

### Materials availability

This study did not generate new unique reagents.

### Data and code availability

The accession numbers for all sequencing data discussed in this manuscript are available in Table S1. Gene expression data have been uploaded to the NIH GEO database under BioProject PRJNA1432846.

## Acknowledgements

We thank the Quantitative Molecular Diagnostics Core of the AIDS and Cancer Virus Program, Frederick National Laboratory, for plasma and cell/tissue viral load analyses.

## METHODS

### Animal Subjects

All study macaques were housed at the Oregon National Primate Research Center (ONPRC) in animal biosafety level 2 rooms with autonomously controlled temperature, humidity, and lighting. Macaques were fed commercially prepared primate chow twice daily and received supplemental fresh fruit or vegetables daily. Fresh, potable water was provided via automatic water systems. During all protocol time points, body weight and complete blood counts were collected, and animals underwent anesthesia support and monitoring. The ONPRC Institutional Animal Care and Use Committee approved macaque care and all experimental protocols and procedures. The ONPRC is a Category I facility. The American Association for Accreditation of Laboratory Animal Care fully accredits the Laboratory Animal Care and Use Program at the ONPRC. It has an approved assurance (no. A3304-01) for the care and use of animals on file with the National Institutes of Health Office for Protection from Research Risks. The Institutional Animal Care and Use Committee adhere to national guidelines established in the Animal Welfare Act (7 U.S. Code, sections 2131-2159) and the Guide for the Care and Use of Laboratory Animals, Eighth Edition, as mandated by the U.S. Public Health Service Policy. Prior to study initiation/infection, all subjects received a multimodal therapeutic regimen to eliminate common gastrointestinal pathogens as previously described^55^. RM were inoculated intravenously with 5000 infectious units of barcoded SIVmac239M^24,56^ before starting daily ART as previously described^23^ [subcutaneous injections of 5.1 mg/kg/d tenofovir disoproxil, 40 mg/kg/d emtricitabine, and 2.5 mg/kg/d dolutegravir in a solution containing 15% (v/v) kleptose at pH 4.2] 9 days post-infection through at least 70 weeks post-infection. The viral dose and timing of ART start was calibrated based on past data in the model to provide a saturated reservoir as well as high resolving power for identifying individual barcode reactivation events, while not impacting acute infection viral dynamics^24,56^.

### Plasma and Cell-associated Viral Load Measurements

Levels of plasma SIV RNA and vDNA and vRNA from cell pellets/tissues were determined essentially as described previously using assays targeting a gag amplicon^22,57,58^.

### Tissue Collection and Processing

Cell isolation from PBMC and solid tissues were acquired and processed to single-cell suspensions using previously published methods, summarized below^59,60^. Spleen and mesenteric lymph node biopsies were collected by a minimally invasive laparoscopic procedure^61^. Bone marrow cells were harvested from the humerus or iliac crest by flushing with R10 media. Peripheral blood mononuclear cells (PBMC) were isolated from freshly collected ACD-A treated blood utilizing Ficoll-Paque density centrifugation (GE Healthcare). Lymph nodes (LN) and spleen samples were homogenized as previously described^62^. Prior to processing, cells were filtered using 70 um strainers. Cells were quantified using a Countess II (Thermo Fisher), aliquoted, diluted as required for single-cell RNA sequencing (typically 500-1,500 cells/uL), and kept on ice prior to processing.

### Cell Hashing

Cell hashing was used for most scRNA-seq samples, with the MULTI-Seq lipid labeling system^63^, using commercially available lipid modified oligos (Sigma Aldridge LMO001). Cells were labeled with barcoded lipids as follows: 15uL MultiSeq solution 1 (LMO stock, diluted in PBS to 400nM) was added, along with 45 uL of the barcode solution (10uM barcode oligo, diluted in PBS to 400nM), giving a final working concentration of 200nM for LMO and 200nM for the barcode oligo. Next, pipet mix and incubate for 5 min at 4°C. Add 10uL of the MultiSeq co-anchor solution (50uM Co-A stock, diluted in PBS to 2uM), then pipet mix and incubate for 5 min at 4°C. Wash twice with 1 mL cold PBS, spinning cells at 700 g for 5 min at 4°C, and then resuspend in 200 uL R10 (which will quench LMOs). Samples were pooled, followed by GEM generation on the 10x instrument.

### Single-Cell RNA Sequencing

The isolated single cell suspensions were then processed for single-cell RNA sequencing using the 10x Genomics Chromium platform, using 5’ v2 or HT chemistry, following the manufacturer’s protocols, including feature barcoding library preparation. To improve capture of MULTI-Seq fragments, we added the following primer, 5’-CCTTGGCACCCGAGAATTCC-3’, at 2.5uM to the 10x cDNA synthesis step. Generation of VDJ enriched libraries followed manufacturer’s instructions with the exception that macaque-specific TCR constant region primers were used in place of human-specific TCR enrichment primers for macaque cells^26^. Primer pairs were used to amplify the alpha, beta, delta, and gamma TCR chains. The concentration of the alpha constant region primer was increased relative to the beta primer to improve amplification. PCR conditions for both reactions were as follows: lid temp 105°C, 98°C 0:45, 12 cycles of: 98°C 0:20, 60°C 0:30, 72°C 1:00, followed by 72°C 1:00 and 4°C hold. Sequence libraries were sequenced using Illumina chemistry, on either Novaseq or HiSeq instruments (Illumina).

### Single-Cell RNA-seq Pre-processing and QC

Raw sequence reads were processed using 10X Genomics Cell Ranger software (version 8.0.1). The resulting sequence data were aligned to the MMul_10 genome (assembly ID: GCF_003339765.1) with NCBI gene build 103. Cell demultiplexing used a combination of algorithms, including GMM-demux, demuxEM and BFF, implemented using the cellhashR package^64–66^. Droplets identified as doublets (i.e. the collision of distinct sample barcodes) were removed from downstream analyses. We additionally performed doublet detection using DoubletFinder and removed doublets from downstream analysis^67^. Data were otherwise processed as previously described^26^. Analyses utilized the Seurat R package, version 5.4^68^. Multiple scRNA-seq datasets are used, including many previously published datasets. A complete listing of the SRA accession numbers for datasets generated for this manuscript are available in Table S1, and under NIH BioProject PRJNA1432846, with count data under GEO accession GSE325004. The Rhesus Macaque Immune Atlas (RIRA) dataset was used for multiple analyses^26^. Clustering and dimensionality reduction used a Seurat-based workflow implemented in the CellMembrane R package. Cells were classified into broad immune cell types using celltypist models from the RIRA resource, and finer cell type annotations were manually assigned based on RIRA models and cluster-specific marker gene expression^26^.

### Plasma Proteomics

NULISA assays were performed at Alamar Biosciences as described previously^69^. Briefly, plasma samples stored at -80°C were thawed on ice and centrifuged at 2,200g for 10mins. 25uL supernatant samples were plated in 96-well plates and analyzed with Alamar’s Inflammation Panel 250 targeting mostly inflammation and immune response-related cytokines and chemokines. The ARGO™ HT platform was used to perform the NULISA workflow, starting with immunocomplex formation with DNA-barcoded capture and detection antibodies, followed by capturing and washing the immunocomplexes on paramagnetic oligo-dT beads, then releasing the immunocomplexes into a low-salt buffer, which were captured and washed on streptavidin beads. Finally, the proximal ends of the DNA strands on each immunocomplex were ligated to generate a DNA reporter molecule containing both target-specific and sample-specific barcodes. DNA reporter molecules were pooled and amplified by PCR, purified, and sequenced using Illumina chemistry. Raw data are available in Table S6A.

### Generation of Pseudo-bulk Profiles and Differential Gene Expression Analysis

Single-cell UMI counts were aggregated into pseudo-bulk expression profiles per sample and broad immune cell type by summing raw counts across cells of the same cell type within each sample. Clustering and dimensionality reduction were performed using the same workflow applied to the single-cell data. For each subject within each tissue and immune cell type, longitudinal shifts were quantified in PCA space by calculating the Euclidean distance between baseline and the corresponding post-infection timepoint using the PC1 and PC2 coordinates. When multiple samples were present for a given timepoint, PC1 and PC2 values were averaged within timepoint prior to distance calculation. This PCA distance provides a summary of the magnitude of transcriptional change over time within the same individual.

To detect DEGs while avoiding false positives, differential gene expression testing was performed on the pseudo-bulk counts using edgeR v4.0.14^25^ workflow implemented in CellMembrane, within each tissue and immune cell type using analysis-specific contrasts. To limit the influence of cells with exponentially more RNA on sample-level aggregation, we applied a cellular RNA abundance-based stratification step, prior to pseudo-bulking, to identify clusters with abnormally high transcript capture, typically corresponding to cycling cells. All-versus-all Kullback-Leibler (KL) divergences were computed between sampled cluster-level RNA abundance distributions, and clusters with summed divergence greater than 2 standard deviations above the mean were excluded before aggregation. P-values were adjusted for multiple testing using the Benjamini-Hochberg (BH) procedure, and genes with FDR < 0.05 were considered differentially expressed and are reported in the supplemental tables.

### sPLS-DA Classification of Pseudo-bulk Profiles

Sparse partial least squares discriminant analysis (sPLS-DA) was used to classify pseudo-bulk profiles by timepoint across tissues for each broad immune cell type. Models were fit in R using mixOmics v6.24^37^. For each cell type, scaled gene expression for the top 5000 variable features were used as input and two components were fit. The number of selected genes per component (*keepX;* 5 to 50 features) was tuned using repeated M-fold cross-validation, optimizing balanced error rate (BER). Final models were refit using the selected *keepX*, and sample scores and gene loadings from the first two components were used for visualization and interpretation.

### Heatmap Visualization, Clustering and GO Annotation

Heatmaps were generated using ComplexHeatmap v2.16^70^ to visualize feature patterns across groups. Values were z-scored within each feature across samples/groups prior to visualization to emphasize relative differences. Features were grouped using k-means clustering, and columns (or rows) were ordered by the relevant experimental grouping with associated metadata shown as annotations. Representative Gene Ontology Biological Process (GO:BP) terms for gene clusters were identified using the *enrichGO* function in clusterProfiler v4.8.3^71^. P-values were adjusted for multiple testing using the BH method and GO:BP terms with adjusted p < 0.05 were considered significant. Redundant GO:BP terms were reduced based on semantic similarity (cutoff = 0.7).

### Gene Module Scoring and Proportion Analysis

Cells were scored for pre-defined gene modules (Table S7) using the UCell v2.12^40^. Module scores were summarized as mean values per sample and immune cell type (or per sample, where indicated). For each group of interest, we also quantified the proportion of cells with module scores exceeding a selected threshold, defined based on the UCell score distributions. Statistical differences in module-positive cell proportions between groups were assessed using the Mann-Whitney U test, with p-values adjusted for multiple testing using the BH method.

### Prediction of T Cell Activation

T cells were classified using the *PredictTcellActivation* function from the RIRA package^26^, which applies a probabilistic model to identify TCR-stimulated T cells. Predicted activation labels were then used to quantify the proportion of activated T cells.

### Cell-associated Viral Load and Interferon Response Population Modeling

Proportions of cellular populations were modeled conditionally using beta regression in brms^72^ to assess the relationship between cell-associated viral load and tissue-level interferon responses. To deal with heteroskedastic responses across tissues, both the mean and precision were conditioned on tissue. Specifically for observations *i* and subjects *j*:

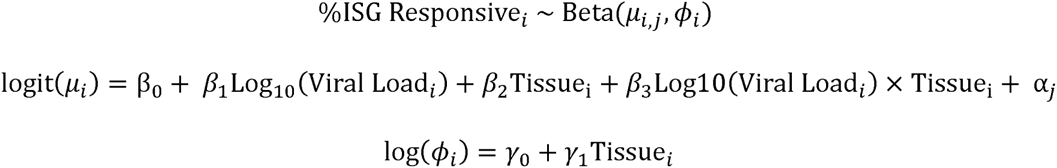

The three point estimates (5^th^ quantile, median, and 95^th^ quantile) reported are Hamiltonian Monte Carlo^73^ estimates of the posterior distributions for the beta’s mean and precision parameters. The default prior parameterization was used. Specifically, the priors were uninformative (i.e. uniform/flat) for the mean/precision parameters *β*_l,2,3_ and *γ*, while a Student-t distribution with three degrees of freedom, zero mean, and 2.5 scale were used for the fixed intercepts (*β*_0_, *γ*_0_) and random intercept (*α*). A small scalar (10^-4^) was added/subtracted from the response variables to shift them into the open interval (0,1) for the beta regression.

### Plasma Proteomic Modeling

The plasma proteomic/NULISA data were analyzed using two linear mixed-effects models^74^. Both models fit non-centered random intercepts to address substantial subject-level clustering. The first model addresses differential abundance (for observations *i* in subjects *j*) between the baseline and acute timepoints via a likelihood ratio test between the two nested models:

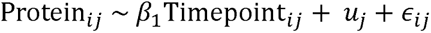

and a timepoint naïve model, containing only a random intercept conditioned on subject ID:

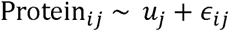

where the random intercepts *u* and errors *ɛ* are assumed normally distributed. The p-values were adjusted via the BH procedure for each tested protein, and those with FDR < 0.05 were defined as differentially abundant. To show the variation of protein abundance over different timepoints, we used z-scored, subject-adjusted, residuals from the timepoint naïve model.

### Pathway Activity Inference

Pathway activity was inferred using decoupleR package v2.9.7^75^ with PROGENy, a curated collection of 14 signaling pathways and their downstream target genes, with weighted gene-pathway interactions^48^. For each tissue and immune cell type, pseudo-bulk differential expression was performed with DESeq2 v1.40.2^76^ by comparing each post-infection timepoint to baseline (or HIV-1 seronegative controls for the human dataset). The RNA abundance-based stratification step was not applied before aggregation for this analysis as we aimed to preserve transcriptional signals related to cell cycling. The resulting gene-level Wald test statistics were used as input to estimate PROGENy pathway activities. Inference was restricted to the top 500 target genes per pathway, ranked by PROGENy p-value, and pathway activity scores were obtained by fitting a multivariate linear model (using *run_mlm* function in decoupleR), for pathways with at least 5 mapped target genes. For visualization, pathway scores were zero-centered and scaled across groups (timepoints/cohorts) within each pathway, tissue, and immune cell type to preserve directionality while normalizing magnitude for cross-group comparisons.

