## Supplemental Figures for "Distinct phases of immune system programming during ART-suppressed immunodeficiency virus infection"

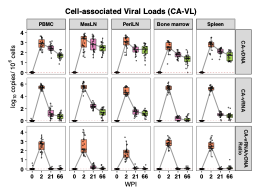

**Figure S1. Cell-associated viral measures in the SIV/ART cohort.** CA-vDNA, CA-vRNA, and CA-vRNA/vDNA ratio measured across PBMC and lymphoid tissues over time in the primary cohort. Red dashed line indicates the assay limit of detection (LOD; 1 copy/10^6^ cells).

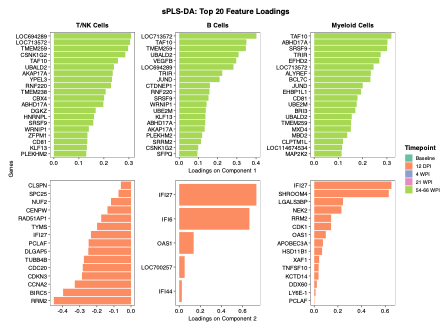

**Figure S2. sPLS-DA feature loadings across immune cell types.** Sparse partial least squares-discriminant analysis (sPLS-DA) identifies genes that best separate longitudinal timepoints in the SIV/ART cohort. Bar plots show the top 20 genes with the largest absolute loadings for Component 1 (top) and Component 2 (bottom) within T/NK, B, and myeloid cells. Genes are ordered by loading magnitude; bar length reflects each gene’s contribution to the corresponding component. Colors indicate the timepoint with the highest mean expression for a given gene.

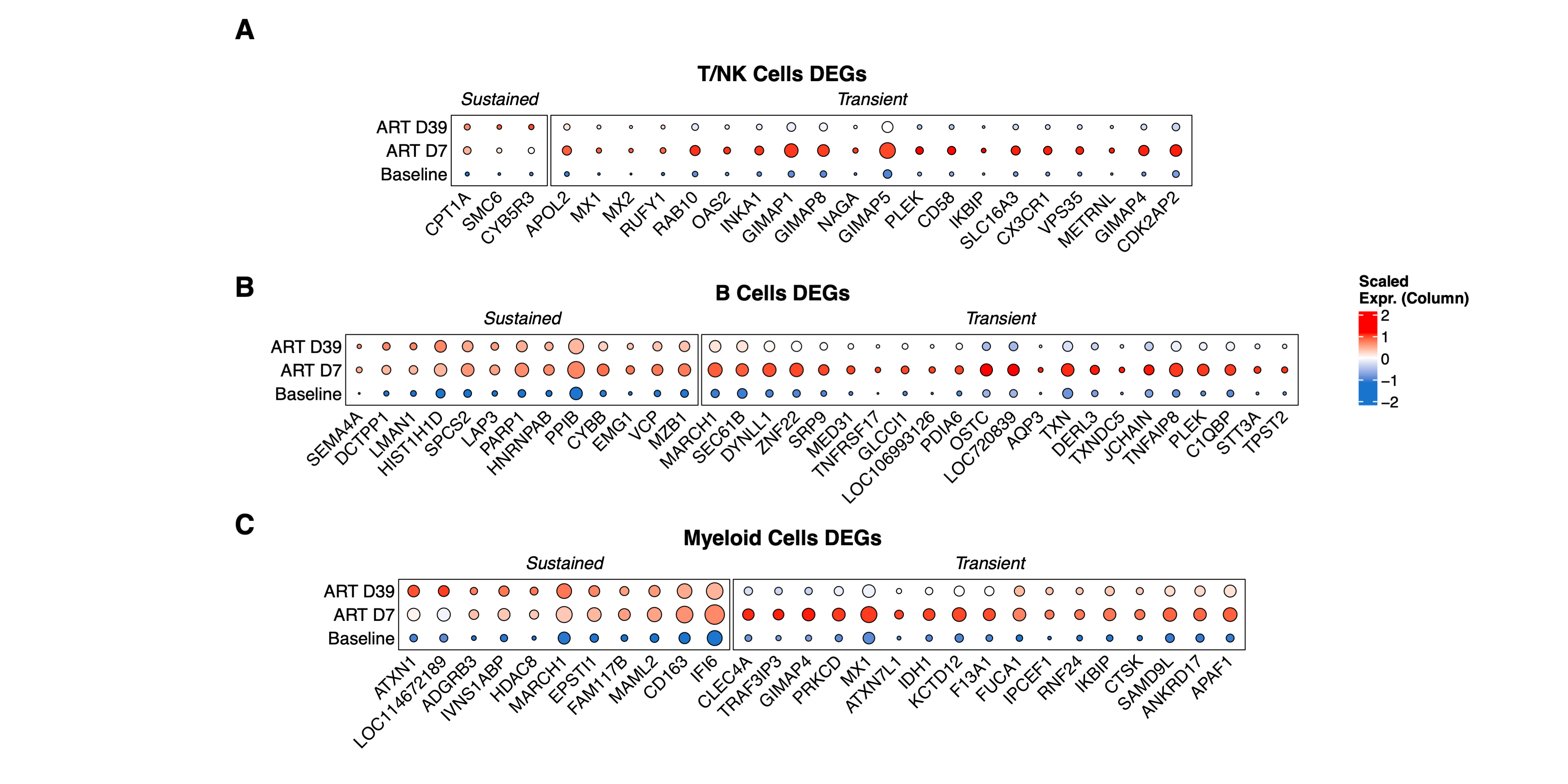

**Figure S3. Effects of ART in the absence of SIV. A)** Dot plots showing top differentially expressed genes (log_2_FC > 1, FDR < 0.05) at ART D7 and ART D39 versus Baseline in the uninfected ART-treated cohort, for the indicated cell types. While most ART-related transcriptional changes are transient, some effects remain as late as D39.

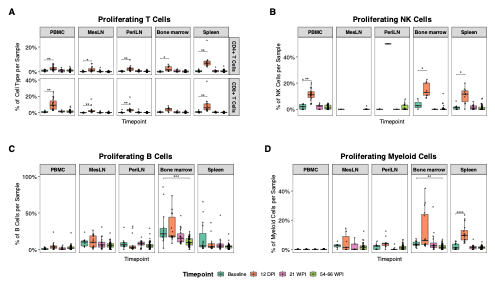

**Figure S4. Quantification of cell proliferation.** All plots summarize single-cell RNA-seq data from the primary SIV/ART cohort. Percentages are within-sample: **A)** of CD4⁺ or CD8⁺ T cells, as indicated; **B-D)** of the indicated lineage. **A)** Percentages of proliferating CD4⁺ and CD8⁺ T cells across tissues and timepoints. **B)** Percentages of proliferating NK cells across tissues and timepoints. **C)** Percentages of proliferating B cells across tissues and timepoints. **D)** Percentages of proliferating myeloid cells across tissues and timepoints.

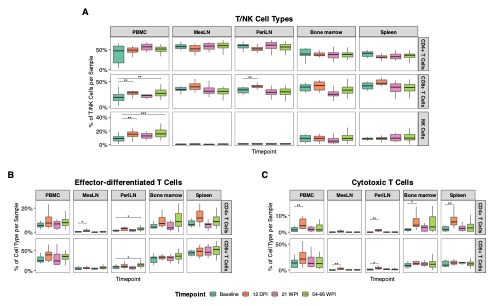

**Figure S5. Quantification of T and NK cell subsets****.** All plots summarize single-cell RNA-seq data from the primary SIV/ART cohort. Percentages are calculated within-sample. **A)** of total T/NK cells; **B-C)** of CD4⁺ or CD8⁺ T cells, as indicated. **A)** Percentages of CD4^+^ T, CD8^+^ T and NK cells across tissues and timepoints. **B)** Percentages of effector-differentiated CD4^+^ and CD8^+^ T cells, defined as T cells with an effector differentiation score (EDS) > 6. EDS, defined in the RIRA resource, was trained on reference peripheral T cells and quantifies the naïve-to-effector differentiation across T cell datasets. **C)** Percentage of cytotoxic CD4^+^ and CD8^+^ T cells, defined by scoring cells using a cytotoxicity gene module (Table S7) and thresholding cells on module score > 0.5.

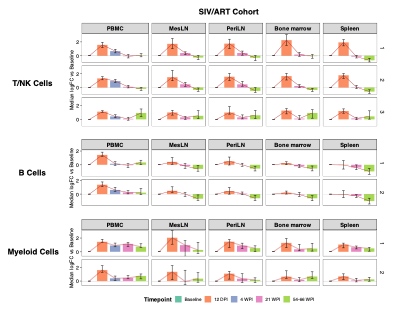

**Figure S6. Tissue-level trajectories for PBMC DEG modules.** Bar plots summarizing the trajectories of the PBMC gene clusters from Fig. 3C for each immune cell type across tissues and timepoints in the primary SIV/ART cohort. For each immune cell type (row; left), cluster (row; right), and tissue (column), gene-level logFC relative to baseline was calculated at each timepoint (**Table S2**) and summarized as the median logFC (bars) with IQR (error bars). Colors indicate timepoints and cluster numbers correspond to those in Fig. 3C

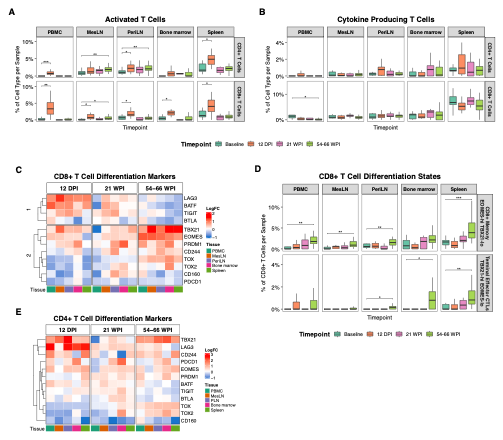

**Figure S7. T cell activation and differentiation.** All panels display single-cell RNA-seq data from the primary SIV/ART cohort. Percentages are within-sample of CD4⁺ or CD8⁺ T cells, as indicated**. A)** Percentages of activated CD4^+^ and CD8^+^ T cells over the indicated tissues and timepoints. Activation is defined by a separately published probabilistic model trained on gene signatures proximal to TCR stimulation. **B)** Percentages of cytokine-producing CD4^+^ and CD8^+^ T cells over the indicated tissues and timepoints. Cytokine production is defined by scoring cells for a cytokine gene module (Table S7), thresholding cells on score > 0.2. **C)** Clustered heatmap of log fold change (logFC; relative to Baseline) in CD8^+^ T cells for selected T cell differentiation / exhaustion-associated genes across tissues and timepoints. **D)** Percentages of two CD8^+^ T cell differentiation states, defined by mutually exclusive transcription factor programs: an EOMES-hi / TBX21-lo memory-like program and an EOMES-lo / TBX21-hi terminal effector/CTL-like program, shown across tissues and timepoints. Differentiation states were defined by scoring cells for the corresponding gene modules (Table S7) and thresholding cells on score >0.6. **E)** Clustered heatmap of logFC (relative to Baseline) for the same gene set as in **(C)**, shown for CD4^+^ T cells across tissues and timepoints.

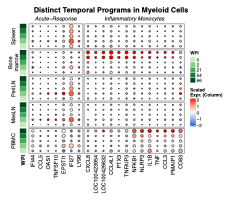

**Figure S8. Inflammatory monocyte gene expression.** The dot plot summarizes gene expression within myeloid cells over the indicated tissues and timepoints. The left-hand module contains genes differentially regulated in acute infection. The right-hand module contains genes enriched in the inflammatory monocyte population that is enriched in bone marrow at 54-66 WPI.

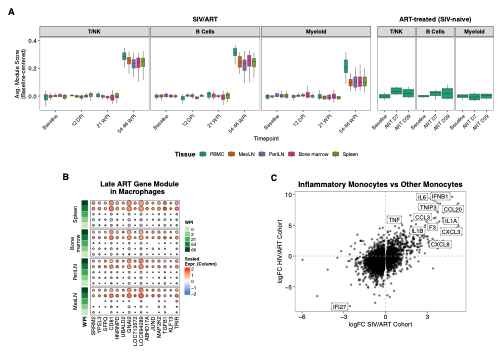

**Figure S9. Late ART Gene Programs across cohorts and cell types. A)** Average Late ART gene module scores (relative to Baseline) across tissues and timepoints for each immune cell type in the SIV/ART cohort and the ART-treated, SIV-naïve cohort. **B)** Dot plot of scaled mean expression of Late ART module genes in macrophages within lymphoid tissues over WPI, showing the induction of this program in macrophages at 54-66 WPI. **C)** Scatter plot of fold change values for shared DEGs comparing inflammatory monocytes with other monocyte clusters in the SIV/ART and human cohorts. Labeled genes highlight a shared inflammatory monocyte signature across cohorts.

**Table S1. List of all raw sequence datasets and NIH SRA accession numbers**

**Table S2. Pseudo-bulk differential expression results for the SIV/ART cohort, comparing each timepoint to baseline within each tissue and immune cell type.**

**Table S3. Pseudo-bulk differential expression results for the ART-treated, SIV-naïve cohort, comparing each timepoint to baseline within each tissue and immune cell type.**

**Table S4. Pseudo-bulk differential expression results for SIV/ART cohort, PBMCs at 12 DPI vs baseline within each immune cell type, including heatmap clusters.**

**Table S5. Pseudo-bulk differential expression results for the SIV/ART cohort, comparing Late ART samples to the combined Pre-infection + Early Infection/ART group within each tissue and immune cell type.**

**Table S6. Alamar plasma proteomics dataset, including all assayed targets, with detectability status indicated.**

**Table S7. List of UCell gene modules used for cell state scoring.**
